# A diversity of traits contributes to salinity tolerance of wild Galapagos tomatoes seedlings

**DOI:** 10.1101/642876

**Authors:** Yveline Pailles, Mariam Awlia, Magdalena Julkowska, Luca Passone, Khadija Zemmouri, Sónia Negrão, Sandra M. Schmöckel, Mark Tester

## Abstract

Traits of modern crops have been heavily selected in agriculture, causing the commercial lines to be more susceptible to harsh conditions, which their wild relatives are naturally better able to withstand. Understanding the developed mechanisms of tolerance present in wild relatives can enhance crop performance under stress. In this study, salinity tolerance traits of two species of wild tomato endemic to the Galapagos Islands, *Solanum cheesmaniae* and *Solanum galapagense*, were investigated. Since these tomatoes grow well despite being constantly splashed with seawater, they could be a valuable genetic resource for improving salinity tolerance in commercial tomatoes. To explore their potential, over 20 traits reflecting plant growth, physiology and ion content were recorded in 67 accessions of *S. cheesmaniae* and *S. galapagense* and two commercial tomato lines of *Solanum lycopersicum.* Salt treatments of 200 mM NaCl were applied for ten days, using supported hydroponics. Great natural variation was evident in the responses of the Galapagos tomatoes to salt stress and they also displayed greater tolerance to salt stress than the commercial lines tested, based on multivariate trait analyses. Although Galapagos tomatoes in general exhibited better tolerance to salt stress than the commercial lines tested, the accessions LA0317, LA1449 and LA1403 showed particularly high salinity tolerance based on growth maintenance under stress. Thus, Galapagos tomatoes should be further explored using forward genetic studies to identify and investigate the genes underlying their high tolerance and be used as a resource for increasing salinity tolerance of commercial tomatoes. The generated data, along with useful analysis tools, have been packaged and made publicly available via an interactive online application (https://github.com/mmjulkowska/La_isla_de_tomato) to facilitate trait selection and the use of Galapagos tomatoes for the development of salt tolerant commercial tomatoes.

## Introduction

High soil salinity is one of the main agricultural challenges in the modern world (Rengasamy, 2016). Salt stress affects the growth and development of plants, thus significantly reducing their yield and productivity (Arzani and Ashraf, 2016). Global cultivated lands cover ~1.5 billion hectares, and an estimated 32 million hectares are damaged by salinity. Irrigated lands, having the highest productivity, comprise just 230 million hectares, of which an estimated 20% have yields significantly reduced by high soil salinity (Munns, 2005). Water availability for agriculture is another major concern, not only in desert regions but at a global level, as freshwater supplies are depleting (Famiglietti, 2014). Salt-affected areas worldwide are predicted to continue expanding at a rate of ~10% per year due to low precipitation, high surface evaporation, erosion of rocks, irrigation with saline water and poor cultural practices (Foolad, 2004).

Wild relatives of modern crops have been used for crop improvement for more than 60 years (Hajjar and Hodgkin, 2007). In their natural habitats, they are able to withstand harsh conditions such as high soil salinity (Muñoz *et al.*, 2017). Adaptation to wide-ranging environments has enriched the genetic pool of these wild relatives (Gruber, 2017), thus representing a rich source of potentially beneficial alleles that can be explored to improve salinity tolerance (Zamani Babgohari *et al.*, 2013). Therefore, performing phenotypic screens to capture the natural variation of the wild germplasm can help uncover new genetic sources for enhancing stress tolerance (Arzani and Ashraf, 2016).

The Galapagos Islands, an isolated environment close to the center of origin of the current domesticated tomato (Blanca *et al.*, 2012), hold a rich genetic diversity of tomato wild relatives, notably the two endemic species, *S. cheesmaniae* and *S. galapagense*, adapted to thrive in harsh environments and highly saline coastal habitats (Rick, 1956; Rush and Epstein, 1976). Previous studies hypothesized that at least some accessions of Galapagos tomatoes, collected from both coastal and inland regions, are able to survive higher NaCl concentrations than the domesticated tomato (Rush and Epstein, 1976, 1981; Tal and Shannon, 1983). However, little is known about the specific mechanisms by which the Galapagos tomatoes thrive on saline soil.

The known mechanisms involved in plant salinity tolerance can be classified into three types: osmotic tolerance, involving the sensing and signalling modules occurring before shoot Na^+^ accumulation and causing reductions in growth rate; ion exclusion, the limitation of ion accumulation in the shoot by ion sequestration in the roots; and tissue tolerance, where high Na^+^ concentrations in the shoot are compartmentalized in the vacuoles to reduce their toxic effects (Munns and Tester, 2008; Roy *et al.*, 2014).

An in-depth characterization of Galapagos tomato accessions would allow to understand the mechanisms for salinity tolerance used within these species, as well as to identify the most tolerant accessions for further studies, thus helping to unlock their potential use as genetic resources for improved salinity tolerance of commercial tomato. However, the wild nature of these plants makes them difficult to compare with domesticated commercial lines, as they differ in growth rates and habit. Hence, developing a robust phenotyping method that is suitable for investigating wild germplasm, accounting for all variations in growth, was necessary. The application of salt treatments at the same growth stage for all accessions is important when growth rates vary significantly. To effectively deliver the salt treatment, a hydroponics growth system is preferred since it allows the precise control of salt concentration in the medium (Genc *et al.*, 2007; Munns *et al.*, 2010; Negrão *et al.*, 2017). Moreover, the effects on the ionic activity of micronutrients, such as calcium, by the interaction between salt and nutrients in the medium, can be calculated and balanced by adding supplemental nutrients (Tester and Davenport, 2003). Also, given so many tomatoes are grown commercially in hydroponic systems, such results can even be of direct relevance to application.

In this study, the phenotypic traits related to salinity tolerance of 67 different accessions of wild tomato from the Galapagos Islands were scored, using a flood-and-drain hydroponic growth system. Traits reflecting growth, physiology and ion content were explored using multivariate analysis, leading to a better and more comprehensive understanding of the role of these traits contributing to salinity tolerance in Galapagos tomatoes. A wide variation in growth, physiology and ion content was observed across the accessions, demonstrating great natural diversity underlying the three main mechanisms of salinity tolerance within the Galapagos tomatoes.

## Materials and Methods

### Plant material and seed treatments

A collection of 67 Galapagos tomato accessions (Pailles *et al.*, 2017) was characterized and screened for salinity tolerance, of which 39 are *S. cheesmaniae* and 28 are *S. galapagense*. Two commercial *S. lycopersicum* varieties were also used for comparison: Heinz 1706 and Moneymaker. Heinz 1706 has a published reference genome sequence (The Tomato Genome Consortium, 2012), and Moneymaker was previously shown to have mild tolerance to salt stress (Cuartero *et al.*, 1992). The seeds were obtained from the Tomato Genetics Resource Center (TGRC), UC Davis, California, USA, and propagated in the greenhouse of King Abdullah University of Science and Technology (KAUST). Seeds from a single plant were used in the experiment.

To sterilize the surface of the seeds and to break their dormancy, all seeds were treated with 10% bleach solution for 10 to 30 minutes, until the seed coat was softened and become transparent, then they were washed several times in tap water. This treatment was necessary for the germination of most of the Galapagos tomato seeds (Rush and Epstein, 1976). Although the commercial varieties did not require bleaching to germinate, the same treatment was applied to all the seeds.

### Experimental hydroponics setup

Eight-centimeter square pots were filled with plastic pellets as a substrate to support the roots. The pellets were chosen for their inert quality and dark color to protect the roots from light. The plastic pellets were made of 20% talc-filled polypropylene, black, and with a density of 1.05 g/cm^3^ to sink in water (Edwards Industrial Repair, Robards, KY). Treated seeds were germinated directly in the pots, on 0.8% agar plugs (8 mm diameter, 12 mm deep) containing ¼ Murashige and Skoog (MS) salts inserted in the plastic pellets (Figure 1A). Two agar plugs, each with one seed, were placed in each pot to increase the chances of germination of at least one seed per pot. Sown pots were placed in nursery trays filled with fresh water and covered with a transparent plastic cover, then kept at 26°C. Germination took between three and eight days. After germination, the two seedlings per pot were thinned to one, by choosing plants with even size and healthy appearance. Treatment with 10% bleach was repeated for those seeds that did not germinate one week after the first treatment (Darwin, 2009). Six biological replicates were used for each control and salt treatment. Because different species with different growth habits were being compared, another six replicates were included, to be harvested before the salt treatment was started – thus, the effects of salinity on growth that occurred only during the time of the salt treatment could be calculated, correcting for differences in growth that occurred prior to the salinity treatment.

**Figure 1:**
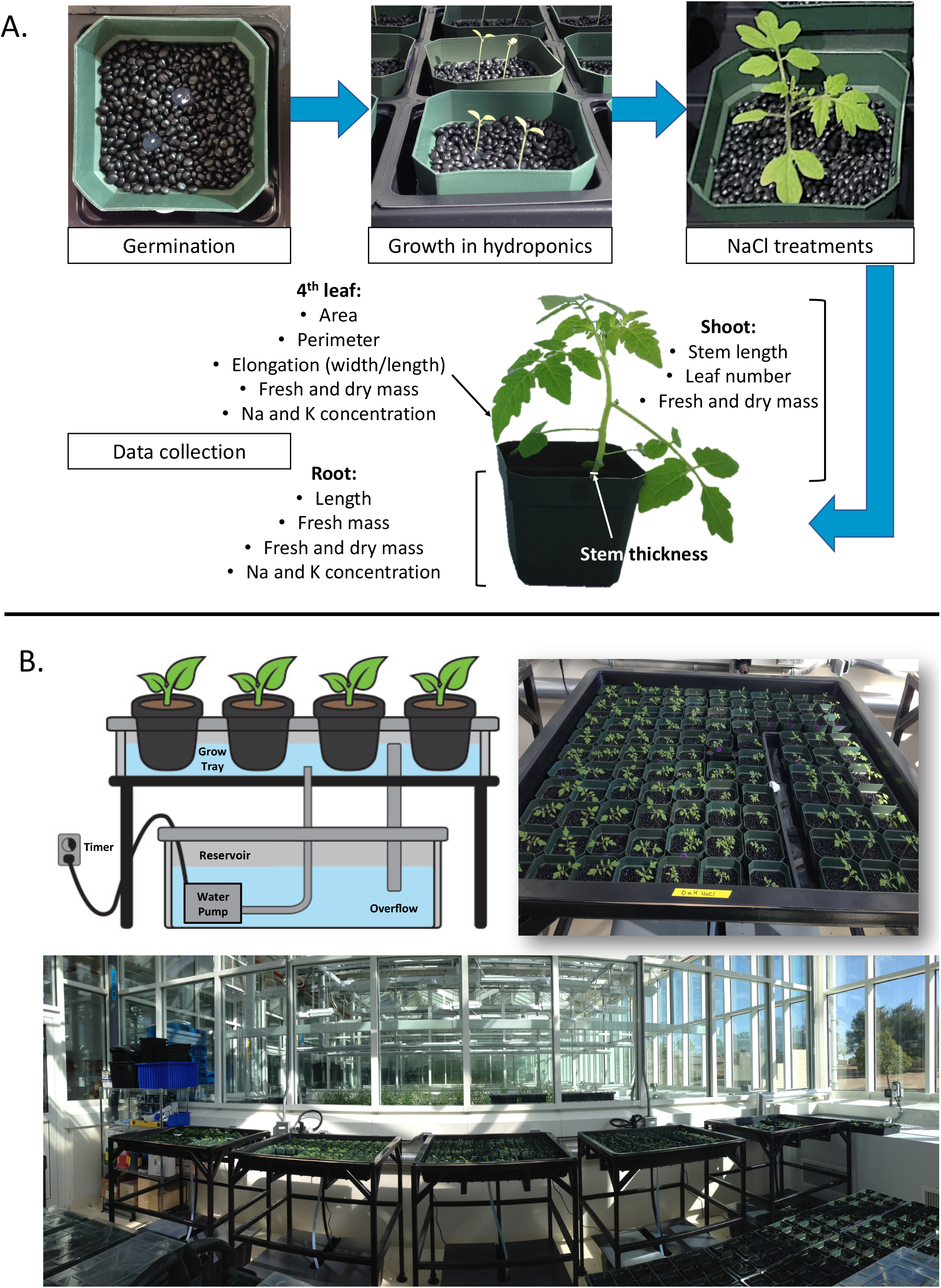
Screening system description. A. Workflow from germination to final harvest: one seed on an agar plug, two agar plugs per pot, pots filled with plastic beads. After germination and thinning to only one plant per pot, they were moved to grow in hydroponics. After 10 days of salt stress, the plants were harvested to record the effects of salinity on their physiology. Recorded traits are listed. B. Supported hydroponics system for screening plants for salinity tolerance at the seedling stage. Diagram adapted from Kruger and Doyle (2016). Pots are held on top of the grow tray. A 100 L reservoir tank with nutrient or saline solution is located under the grow tray and a pump is used to feed the solution up to the grow tray. The pump is controlled by a timer, programmed for 15 min ON/OFF intervals. Solution floods the grow tray for 15 minutes and drains back into the reservoir tank when the pump turns off for another 15 min.

When the cotyledons had emerged fully and the radicle was long enough, the pots were transferred from the nursery to a supported hydroponics system (EconoTray by American Hydroponics, Inc.), which uses an ebb-and-flow scheme for root aeration (Figure 1B). This system consists of a grow tray, a tray frame, a 100 L nutrient reservoir tank, and a submersible aquarium-pump (ViaAqua 360 by Commodity Axis, Inc.). The tray frame height was modified, from 0.5 to 1.06 m, to hold the plants above the greenhouse walls and thus avoid shading. Each growth tray was able to hold up to 108 pots (8 cm^2^) with seedlings at the cotyledon stage and 96 pots (8 cm^2^) growing tomato plants up to the 7^th^ or 8^th^ leaf stages. The growth tray rests on the tray frame above the reservoir tank, which contains the nutrient solution. The solution was pumped to the grow tray to deliver nutrients to the plants, then drained back into the reservoir tank allowing root aeration. Aquarium pumps inside the reservoir tanks were controlled by a programmed timer to be on for 15 min, pumping nutrient solution up to fill the grow tray, and off for 15 minutes, allowing the solution from the grow tray to drain back into the tank. The nutrient solutions were prepared using 100 L of tap water from the greenhouse and 33 ml of each nutrient stock, FloraGro, FloraMicro and FloraBloom (General Hydroponics), as suggested by the manufacturer. The tap water was tested for calcium, chloride, potassium, sodium, ammonium and nitrate ion content using Multi-Ion Kit (CleanGrow Europe), before preparing the solutions. These measurements were recorded for future normalization. Nutrient depletion was monitored weekly using the Multi-Ion Kit ion-sensitive electrodes (CleanGrow Europe). The nutrient solution was changed at the start of the salt stress treatment and no further significant nutrient depletion occurred throughout the rest of the experiment (10 days).

### Salt stress treatment and considerations

The use of 8 units of the hydroponics system allowed the salt treatment of a total of 69 accessions of three different tomato species (*S. cheesmaniae, S. galapagense*, and *S. lycopersicum*), with 6 biological replicates per accession per treatment, plus 6 seedlings per accession that were harvested as a baseline before treatment. The plants were subjected to salt stress treatment at the same developmental stage, when the 4^th^ leaf started to emerge. Given that *S. lycopersicum* plants were bigger than Galapagos tomatoes throughout all developmental stages (as leaves were larger and stems were thicker), they were treated when the 3^rd^ leaf started to emerge, to compensate for the size difference (Figure S1), since bigger plants are often better able to tolerate salt stress than smaller plants.

The salt stress treatment was administered gradually to the plants. As each plant reached the desired developmental stage (i.e. the 4^th^ or 3^rd^ leaf emergence), it was moved from the nutrient-only hydroponics systems to the systems supplemented with NaCl. Plants were first moved to a 75 mM NaCl hydroponic system and 12 hours later, moved to a 200 mM NaCl hydroponic system. Seedlings remained in the 200 mM NaCl hydroponic system for 10 days. Supplemental CaCl_2_ was added to the solutions to compensate for the decrease in Ca^2+^ activity arising from the addition of NaCl (Tester and Davenport, 2003). The amount of CaCl_2_ added to the NaCl solutions was calculated using GEOCHEM-EZ software (Shaff *et al.*, 2010) to maintain Ca^2+^ activity at 0.4 mM, which was the normal Ca^2+^ activity in the nutrient solution prior to NaCl addition (Table S1).

### Sample collection and recording traits related to salinity tolerance

Plants were photographed and tissues harvested to measure traits related to plant growth, leaf area, and ion allocation (Figure 1A). Photographs of each plant were taken at the start and end of the salt treatment, using a Photosimile 200 light-box and a Nikon D5100 digital single-lens reflex camera. The in-camera white balance calibration function was used with a reference photo of the empty light-box white background, taken at the intended light intensity. This ensured a consistent and accurate colour capture in all photographs taken. The images were used to test a non-destructive approach to estimate the salinity tolerance of Galapagos tomato seedlings. The photographs were processed using a Matlab script for green pixel count (green_finder_V2.m), which can be found in the Supplementary Data. For destructive sample harvesting before and after salt treatment, the plant was carefully extracted from the pot, roots were rinsed in 10 mM MgCl_2_ solution with the excess solution dried off using tissue paper. Plant root, shoot, and 3^rd^ or 4^th^ leaf tissues were each weighted separately. The 3^rd^ and 4^th^ leaves, taken from *S. lycopersicum* and Galapagos accessions respectively, were further characterized as they had developed under salt stress conditions. Each leaf was scanned using an EPSON scanner to calculate leaf area, perimeter, height and width using the WinFolia software (Regent Instruments Inc.). In terms of physical measurements, stem thickness was measured at the base using a caliper, while the stem and root length were measured using a ruler. The different tissues were then stored in paper envelopes and dried at 60**°**C for three days to then measure their dry mass. Dry leaf samples (without petiole) and root samples were digested in 50 mL Falcon tubes with 5 mL of 1% (v/v) nitric acid in a HotBlock™ (Environmental Express) at 80**°**C for four hours. Sodium content was measured in leaf and root samples using a flame photometer (model 420; Sherwood Scientific Ltd., Cambridge, UK).

### Data analysis

A multivariate analysis was performed to assess the effect of the salt-induced changes in the plants under salt stress conditions. The mean of six replicates was calculated for all the measured traits, with the exception of leaf number, for which the mode was calculated instead.

All traits were corrected by subtracting the initial measurement (before salt stress treatment) to measure only the differences that occurred during treatment (Figure S2), except for traits measured in leaves 3 or 4. To facilitate comparison of different species and accessions with large differences in early growth (Figure S3), only traits of salt-treated plants relative to traits of control treated plants were used for analysis (Negrão *et al.*, 2017):

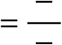

To determine the salt stress effects on the different accessions, correlations between all of the traits measured were analysed using the *corrplot* package (Wei and Simko, 2016) in R (R Core Team, 2017). A total of 11 representative traits were selected for further analyses.

The variability in traits related to salinity tolerance was described using a principal component analysis (PCA) on a matrix of 11 traits × 64 Galapagos and commercial tomato accessions. Note that of the 67 accessions screened, five had no survivors to the salt stress treatment: LA0526, LA0930, LA1141, LA1411, and LA1815. Since the variables have different units, they were scaled to have a variance of 1 and a mean of 0, by subtracting the mean and dividing by the standard deviation, using the *scale* function in R. PCA analysis was carried out using the *FactoMineR* package (Lê *et al.*, 2008) in R (R Core Team, 2017).

The two Galapagos tomato species were observed to have distinctly different morphologies, hence the phenotypic data for each species were analyzed separately. The K-means clustering method (MacQueen, 1967) was used to reveal groups within the data. The clustering method was run with different numbers of clusters (2 to 6) and it was found that two clusters provided the most interpretable output, in terms of accession clustering by traits. The accessions were grouped based on 11 traits related to salinity tolerance: Na and K concentration in root and leaf, leaf area, leaf elongation (width/length), leaf succulence (leaf water/leaf area), leaf number, stem and root length, and total fresh mass. K-means were calculated using the *stats* package in R (R Core Team, 2017).

To identify possible tolerance mechanisms, all trait measurements from the accessions of each species were compared using a heat map, drawn by the function *heatplot* of the *made4* R package (Culhane *et al.*, 2005), which also draws dendrograms of the traits and accessions using correlation similarity metric and average linkage hierarchical clustering (Eisen *et al.*, 1998).

All the phenotypic data was integrated into an Isla_Tomate App, available at https://mmjulkowska.shinyapps.io/La_isla_de_tomato/. The App allows interactive exploration of the correlations between individual traits as well as cluster analysis of the accessions based on the chosen traits. The App was developed with the shinyapp package. The code used for the App is available at https://github.com/mmjulkowska/La_isla_de_tomato, and the instructions on how to use the App can be found at https://mmjulkowska.github.io/La_isla_de_tomato/.

## Results

### Galapagos tomatoes are more salt tolerant than the commercial tomato varieties tested

As suggested by Negrão et al. (2016), responses to salinity stress were measured only for the time when the plants were stressed (by taking measurements before and after the stress treatment). *S. cheesmaniae* and *S. galapagense* accessions were better able to maintain growth (based on dry mass) during the salt stress period than the *S. lycopersicum* varieties tested (Figure 2). This effect is less apparent when biomass is determined only at the endpoint of the experiment (because of the size advantage of *S. lycopersicum* accessions) (Figure S2). There was a large variation in salinity tolerance between accessions, ranging from a difference in dry mass in saline conditions relative to control conditions of 12 to 55% (Figure 2).

**Figure 2:**
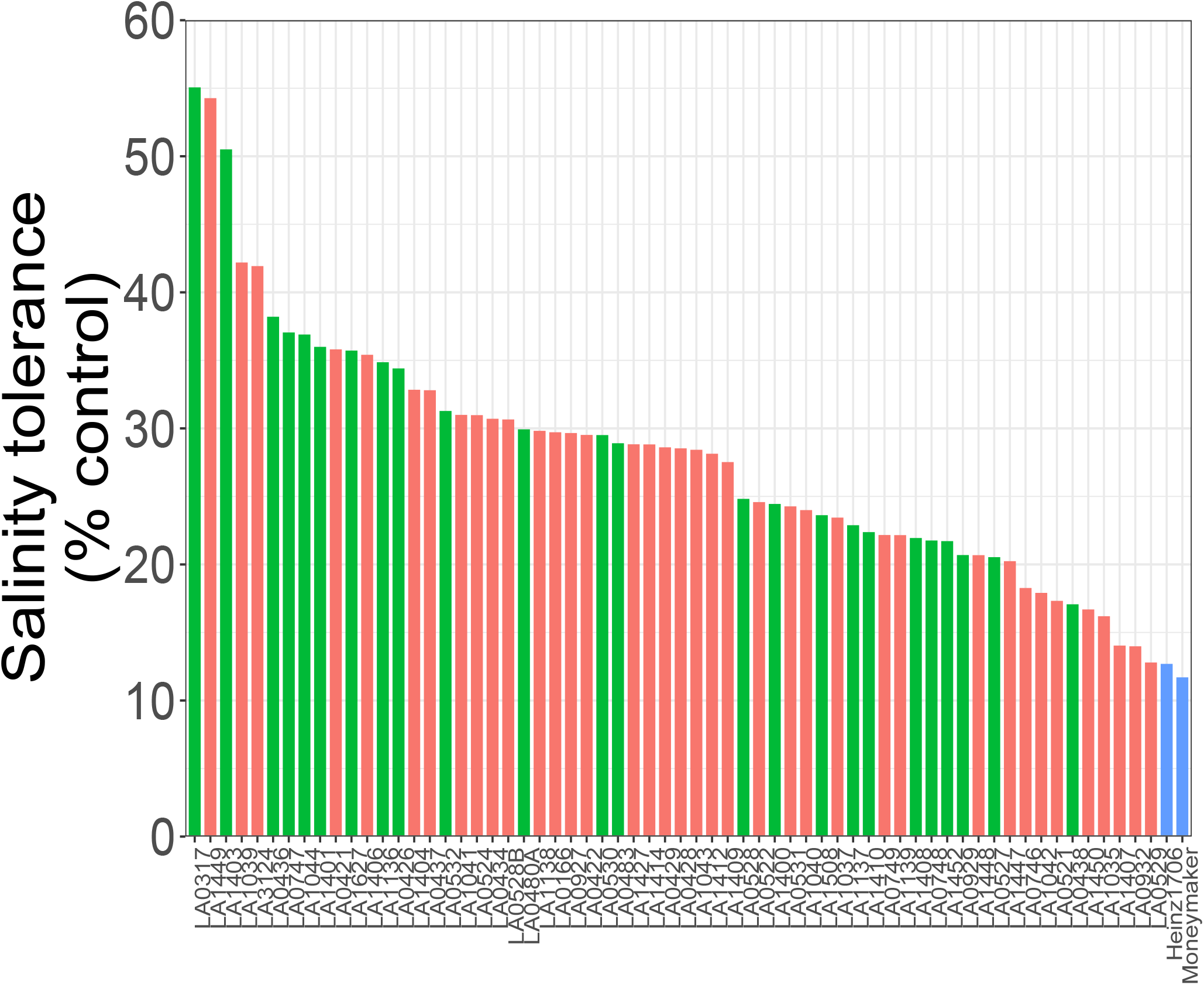
Salinity tolerance across the studied accessions, measured as the difference in dry mass between the start and end of the treatment of plants grown in saline conditions relative to plants grown in control conditions. Different colours represent the different species, *S. cheesmaniae* (coral), *S. galapagense* (green), *S. lycopersicum* (blue).

### Correlation analysis of different seedling traits revealed trait groups

The correlation matrix (Figure 3) shows that leaf traits such as perimeter, vertical length, dry and fresh mass, horizontal width and area are all positively and significantly correlated (correlation coefficients 0.79-0.93, *p*-value=0.001: Figure S4). Plant growth-related traits, such as shoot fresh and dry mass, root fresh and dry mass, total fresh and dry mass and total water content, are also positively and significantly correlated (correlation coefficients 0.69-1.00, *p*-value=0.001) (Figure S4).

**Figure 3:**
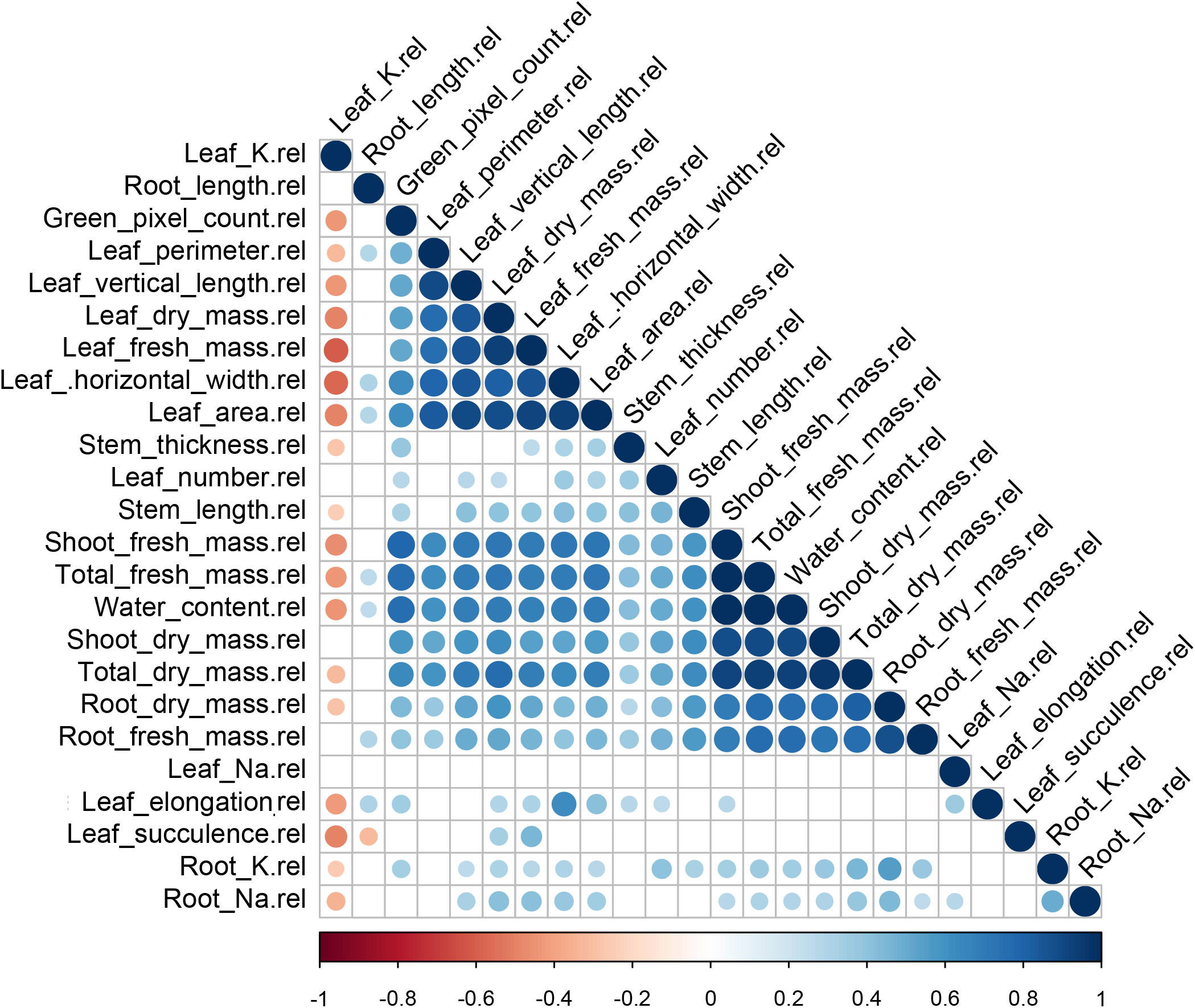
Pearson correlation matrix of the recorded traits in salinity relative to control conditions. Large circles represent strong correlations and small circles represent weak correlations. The color scale bar indicates the sort of correlation, where 1 denotes completely positive correlation (dark blue) and −1 denotes completely negative correlation (dark red) between two traits. Only significant correlations are shown (*p* value<0.05).

Interestingly, leaf K concentration in salt-treated plants relative to control plants is negatively correlated with all the leaf traits, some plant growth-related traits, and Na and K in the root. On the other hand, leaf Na concentration in salt-treated plants, relative to control plants, did not have a significant correlation with any other of the measured traits. In most cases, leaf Na concentration in control plants was negligible, so the leaf Na concentration in salt-treated plants, relative to control plants, was very similar to the leaf Na concentration in salt-treated plants. Na and K in the root have a slight positive correlation with leaf and plant growth-related traits.

Green pixel count was significantly correlated with most of the salinity-tolerance related traits (Figure 3). For most of the accessions tested, the green pixel count had a positive correlation with the shoot fresh mass of salt-treated plants relative to control plants (Figure S5A). However, green pixel count proved a more useful measure of plant growth in *S. cheesmaniae* (r^2^=0.85), than in *S. galapagense* (r^2^=0.64) (Figure S5B-D). Based on the correlation analyses, a set of plant traits representing each of the groups of traits that have been identified as potentially useful predictors of salinity tolerance in plants, were selected for further analyses: Na and K concentration in root and leaf, leaf area, leaf elongation, leaf succulence, leaf number, stem and root length, and total fresh mass. To compare the different species, the traits in salt stress relative to control conditions of the same accession were used.

### Principal component analysis revealed selected traits tendencies and contributions

A PCA was performed to reduce data dimensionality and reveal the potential relationships among representative salinity-tolerance traits. In this study, the four main PCA axes had eigenvalues larger than 1 (Table S2), which indicates that each principal component (PC) accounts for more variance than accounted-for by one of the original variables in the standardized data. This was used as a cut-off to determine the number of PCs to retain.

The PC1 explained 33.8% of the total variability between traits/individuals and was associated with most traits, except leaf Na concentration and leaf succulence (Table 1 and Figure 4). The most significant trait for PC1 was the total fresh mass (Table 1). The accessions at the lower end of PC1 are those whose growth was most affected by salinity but were still able to retain high levels of K in the leaf, while at the higher end, there are the accessions with higher levels of plant growth, leaf area, leaf number, and stem and root length.

**Table 1:** Contributions of plant traits to the four main PCA axes (eigenvalue > 1), obtained from a matrix of 11 traits × 64 Galapagos and commercial tomato accessions. Values are ranked in order of magnitude in PC1. Traits significantly correlated to each PCA axis (α = 0.05) are indicated in bold. All the traits represent the trait value under salt stress relative to control conditions.

| **Traits in salt stress relative to control conditions** | **PC1**<br>**(33.8%)** | **PC2**<br>**(15%)** | **PC3**<br>**(11.7%)** | **PC4**<br>**(9.69%)** |
| --- | --- | --- | --- | --- |
| Plant fresh mass | **18.1** | 1.63 | 3.83 | 0.69 |
| Leaf area | **15.8** | 0.09 | 1.40 | 4.24 |
| Green pixels | **12.7** | **5.43** | **12.7** | 2.36 |
| Stem length | **10.4** | 0.42 | 0.92 | **6.06** |
| Leaf K<sup>+</sup> | **9.76** | **12.5** | **6.83** | **8.04** |
| Root K<sup>+</sup> | **8.12** | 0.39 | 0.54 | **21.9** |
| Leaf number | **8.08** | **4.47** | **8.66** | **12.0** |
| Leaf elongation | **7.77** | 0.45 | **10.0** | **23.3** |
| Root Na<sup>+</sup> | **5.72** | **12.8** | **4.39** | **8.92** |
| Root length | **2.65** | **15.4** | **8.19** | **8.38** |
| Leaf succulence | 0.50 | **35.5** | **5.44** | 0.01 |
| Leaf Na<sup>+</sup> | 0.34 | **10.8** | **37.0** | 3.98 |

**Figure 4:**
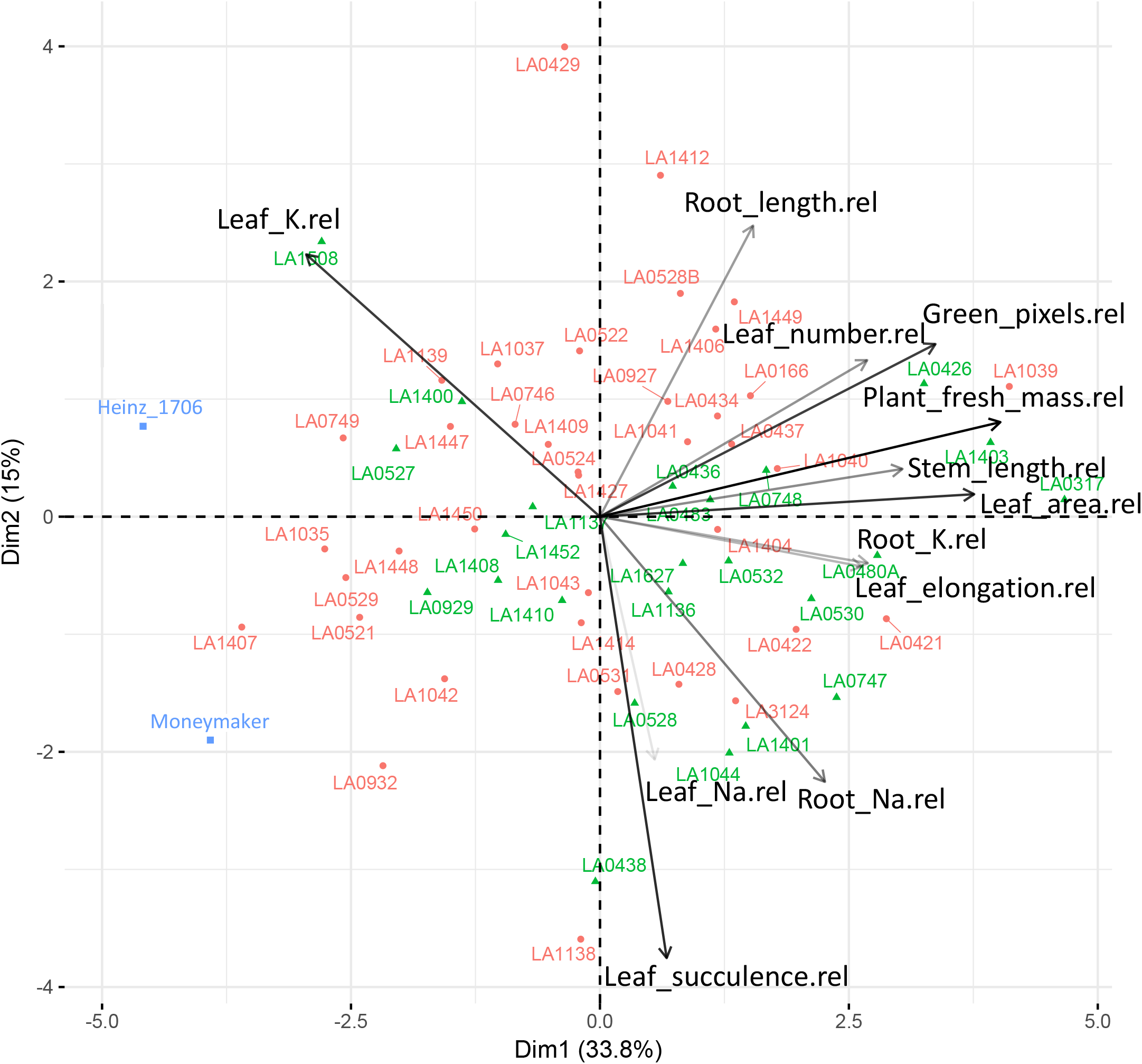
PCA biplot of 62 Galapagos tomato accessions and 2 commercial tomato varieties, based on the variance in 11 salt-stress related physiological traits, explained by two principal component axes. The two components explained 33.8% and 15% of the variance, respectively. Arrows denote the strength of the trait influence on the first two PCs. The transparency of the arrows indicates the contribution to the variance in the dataset, ranging from 5% (lightest) to 12.5% (darkest). The direction and length of the arrows indicate how each trait contributes to the first two components in the PCA. Aligned vectors indicate a strong positive correlation between the two traits. Vectors at right angles/opposites indicate no correlation/negative correlation, respectively. This analysis shows that all growth-related traits are correlated and places the individual accessions where the corresponding. The first component shows that leaf K concentration is negatively correlated with the other representative traits. The second component shows leaf K concentration is negatively correlated with the other representative traits. Individual accessions are placed on the ordination plane. Different colours and symbols represent the different species, *S. cheesmaniae* (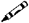 - coral), *S. galapagense* (➡ - green), *S. lycopersicum* (᷋ - blue).

PC2 accounted for an additional 15% of the total variability among seedling traits and appeared to be related to the ion content and some growth traits (Table 1 and Figure 4). The accessions with succulent leaves and higher accumulation of Na in the leaf were located at the lower end of PC2, while those with increased leaf number and K retention in the leaf were located at the higher end of PC2. The PC2 also divided the root Na concentration and root length, where those accessions with high Na concentration in the root had the shortest root.

PC3 accounted for 11.7% of the total variability among salinity tolerance-related traits. It was significantly associated with total fresh mass but had a stronger association with leaf traits, such as elongation factor (length/width) and Na concentration (Table 1). This could suggest that Na concentration in the leaf is independent of the other traits.

PC4 accounted for an additional 10% of the total variability and is significantly associated with Na and K accumulation in the leaf and root, but also, with leaf number and area, and root length (Table 1).

Overall, the PCA indicates that in this experiment, the primary traits that varied and correlated with each other were total fresh mass, ion content and some leaf traits, such as elongation factor.

### Cluster analysis suggests that salinity tolerance at the seedling stage is best defined by the ability of the plant to maintain growth under salt stress conditions

Cluster analysis is a suitable method to study large datasets, involving multiple variables. It allows the grouping of accessions with similar traits and the recognition of hidden patterns or trends in the data. To analyse how the different accessions grouped by traits related to salinity tolerance and to see if any of the traits predominantly explain the overall variation.

The K-means cluster analysis (MacQueen, 1967) of the surviving 38 accessions of *S. cheesmaniae* and 24 accessions of *S. galapagense* after treatment, was used to classify accessions into different clusters (K), where the accessions within the same cluster are as similar as possible, while accessions from different clusters are as dissimilar as possible. The number of clusters K=2 was chosen with the aim of separating the most tolerant accessions from the least tolerant while considering some salinity-tolerance related traits. A total of 11 non-redundant traits were selected, based on the strong correlation of these 11 traits with other traits measured but not among each other (Figure 3).

Considering the values of the selected traits, the Euclidean distance between each accession and the cluster mean was calculated to assign the accession to the nearest cluster. A new mean value of each cluster was calculated after an accession was assigned to it and every accession was checked again to see if they were closer to a different cluster. These steps were iteratively repeated until convergence was achieved.

Bar plots were used to visualize the distribution of the accessions by cluster for each specific trait, a similar visualization strategy is commonly used when plotting Q-matrices and identifying K clusters in population structure studies (Pritchard *et al.*, 2000). The accessions were arranged in descending order and the bars are colored by cluster (Figure S6). By visualizing bar plots for all traits, it was easy to identify that the plant fresh mass was predominantly defining the clustering by K=2. From this, it was observed that the accessions of both species were best grouped by their fresh mass production under salt stress relative to control conditions (Figure S6). Thus, the two clusters divide the accessions of each species of Galapagos tomato into those with high tolerance and low tolerance to salinity, in terms of growth maintenance (Figure 5). Cluster 1 included accessions with higher fresh mass production during salt stress relative to control, indicative of their ability to better maintain growth under salt stress. Cluster 1 of *S. cheesmaniae* had 23 members (Figure 5A) and cluster 1 of *S. galapagense* had 14 members (Figure 5B).

**Figure 5:**
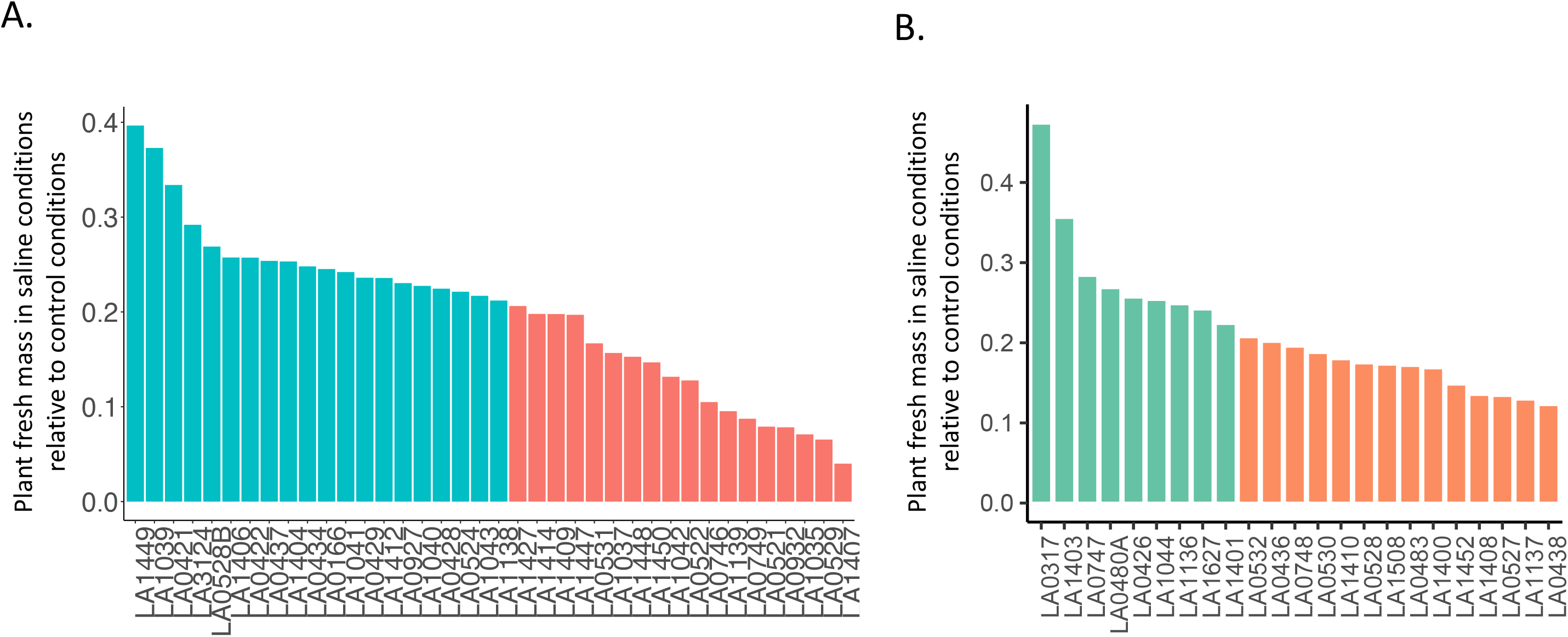
K-means clustering (K=2) of each one of the Galapagos tomato species. **A** barplot for each trait was plotted with the accessions in descending order, different colors represent different cluster assignment. Cluster number was chosen based on the most informative grouping. A. *S. cheesmaniae* within-cluster sum of squares by cluster: 217.6 and 136.9 respectively (between_SS / total_SS = 20.2%). B. *S. galapagense* within-cluster sum of squares by cluster: 115.9 and 94.1 respectively (between_SS / total_SS = 23.9%). Both species showed a clean cluster separation when accessions were arranged in descending order by the total plant fresh mass in salt stress relative to control conditions.

### Natural variation exists across Galapagos tomato accessions in terms of their salinity tolerance mechanisms

Phenotypic data were also analyzed using a hierarchical clustering approach (Figure 6), which was found to be more complex, but more informative, than the K-means clustering method. A heat map paired with the dendrogram obtained by hierarchical clustering, provide a way to visualize and simplify large datasets. This method is routinely used for gene expression data analysis (Eisen *et al.*, 1998) and metabolomics (Tikunov *et al.*, 2005). More recently, it has proven useful also to analyse genotypes and traits interactions (Chen *et al.*, 2014; Julkowska *et al.*, 2016; Clark, 2016; Awlia *et al.*, 2016).

**Figure 6:**
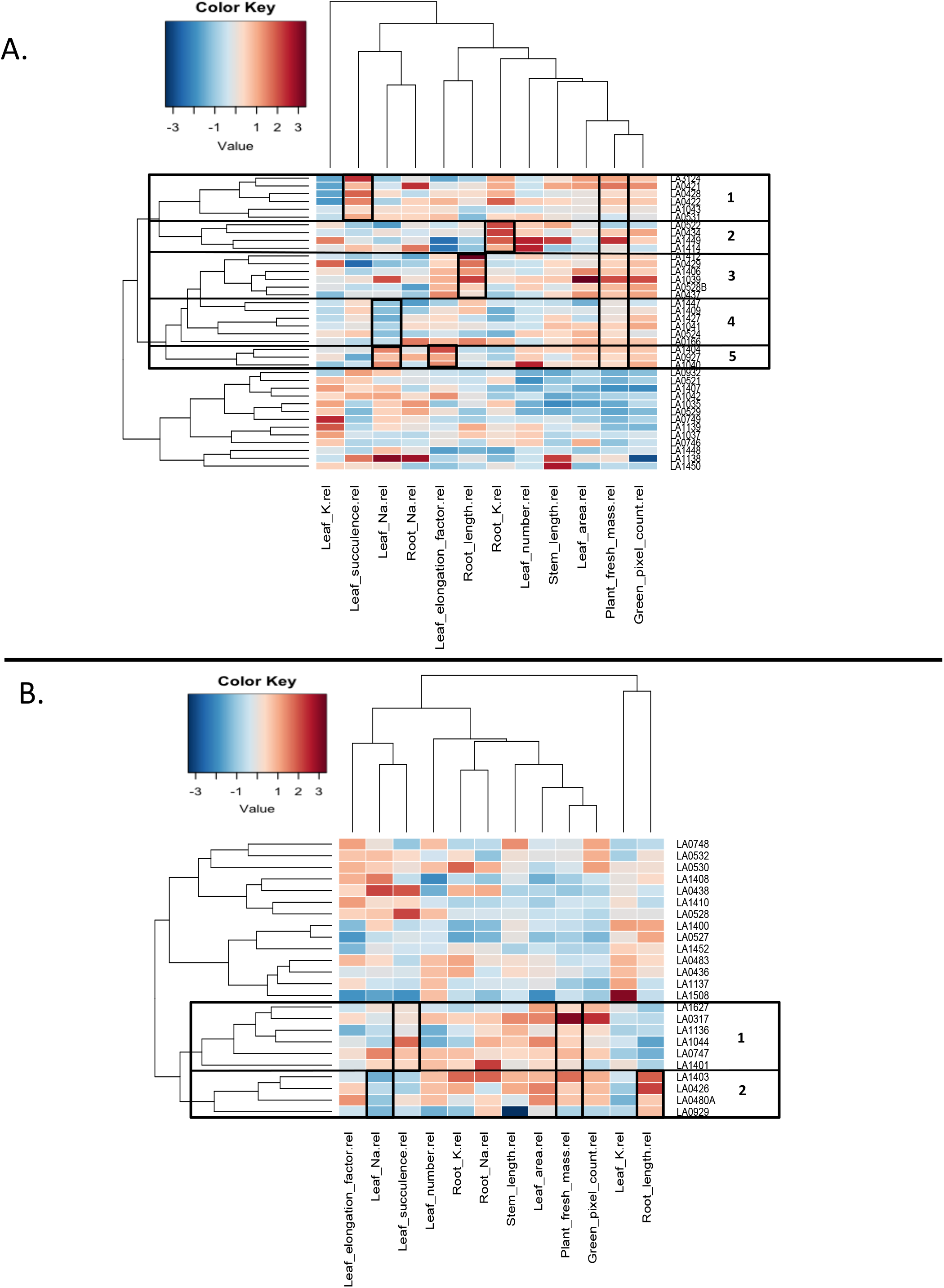
Hierarchical clustering and heat map of all accessions and important salinity-tolerance related traits, divided by species. Each column represents a trait and each row represents an accession. Accessions with large fresh mass, in salt stress relative to control conditions, clustered together. However, they differed in other traits. Further clustering by the similarity in traits is indicated in the figure, showing diversity in salt stress responses among Galapagos tomato accessions. A. *S. cheesmaniae* accessions divide into two main clusters. The accessions in the upper cluster share high values of total fresh mass. This cluster is further divided into 5 other clusters (indicated in the figure) with a distinctive trait that differentiates each cluster from other clusters (Table 2). B. *S. galapagense* accessions divide into two main clusters. The accessions in the lower cluster share high values of total fresh mass. This cluster is further divided into 2 other clusters (indicated in the figure) with a distinctive trait that differentiates each cluster from other clusters (Table 2).

The heat map divided the accessions into two clusters, also defined by fresh mass in salt-treated plants relative to control plants. Moreover, the heat map showed clustering of accessions based on other traits within the main mass-related clusters (i.e. K and Na concentrations in both leaf and root differed greatly between groups of accessions, despite them having similar plant fresh mass). This indicates a pronounced natural variation in salinity tolerance mechanisms within the Galapagos tomato collection.

In *S. cheesmaniae*, the hierarchical clustering method (Figure 6A) separated the accessions into two main clusters, which display contrasting values of fresh mass, leaf area, stem length, leaf number, root length, leaf elongation, leaf Na and K concentration and green pixel count. Within the cluster of accessions with high relative fresh mass, five different clusters could be distinguished, that differed in leaf succulence, root K concentrations, root length, leaf Na^+^ concentration and leaf elongation (Table 2).

**Table 2:** General description of the main mechanisms identified in each of the clusters.

| **Species** | **Highly tolerant accessions cluster** | **High maintenance of particular feature in saline conditions:** |
| --- | --- | --- |
| *S. cheesmaniae* | 1 | Leaf water |
|  | 2 | Root K<sup>+</sup> |
|  | 3 | Root length |
|  | 4 | Leaf Na<sup>+</sup> exclusion |
|  | 5 | Leaf elongation and Na<sup>+</sup> compartmentalization |
| *S. galapagense* | 1 | Leaf water |
|  | 2 | Leaf Na<sup>+</sup> exclusion and root length |

The *S. galapagense* accessions also separated into two clusters (Figure 6B), based on their relative fresh mass, leaf area, root Na, root K, leaf K, and green pixel count. The cluster with high relative fresh mass was divided into two clusters differing in leaf succulence, leaf Na concentration and root length (Table 2).

The phenotypic data collected were integrated into a Shiny App: Isla_Tomate, https://mmjulkowska.shinyapps.io/La_isla_de_tomato/, allowing interactive clustering of the data. The identified clusters can be validated using the Isla_Tomate App, by grouping the accessions into clusters, based on the chosen trait and examining the significant differences between the clusters based on each trait. Significance was calculated using Tukey pair-wise comparison with p-value < 0.05. The data for both species showed that clustering by plant fresh mass forms two significant groups (Figure S7), which we can divide into the two groups of high and low salinity-tolerance accessions.

The dendrograms presented in Figure 6 represent how similar individual accessions react to salt, based on the selected traits. When this grouping of the accessions was compared to their geographical origin or genetic distance between them (Pailles *et al.*, 2017), no correlation was observed. This suggests that geographical origin and evolutionary history do not influence salinity tolerance mechanisms.

The two types of trait association analyses (PCA and clustering analyses) indicated similar trait influences determining differences between accessions and their responses to salinity.

## Discussion

Wild relatives of modern crops surely possess useful traits that can potentially improve plant performance under salt stress. Hence, there is a need for characterizing and screening the available germplasm. However, their wild nature and great natural variation in many traits make them difficult to study quantitatively with conventional methods. The specific mechanisms through which Galapagos tomatoes are tolerant to salinity are not known. Identification of salinity tolerance mechanisms in Galapagos tomatoes will facilitate the improvement of salinity tolerance in current tomato elite varieties.

In this study, most available accessions of wild tomatoes endemic to the Galapagos Islands were screened for salinity tolerance traits. The first objective was to develop an efficient screening method that allowed the quantitative comparison of salt stress responses of different wild tomato seedlings. For this, a commercial ebb-and-flow supported hydroponics system was used. The use of hydroponics for experimental screenings allows better control of the growth media, stress exposure and experimental reproducibility (Genc et al. 2007, Munns et al. 2010). In this case, the hydroponics system facilitated an even delivery of NaCl to the plants at the root level. The plastic-beads substrate conserved moisture during drainage, supported the roots, and protected them from breaking and allowing uncontrolled Na^+^ influx into the root system (Miller, 1987). An opaque substrate was preferred to simulate the light blocking properties of soil and to limit algal growth. The system’s flexibility allowed treatment at the same developmental stage, independent of the growth rate of each of the different genotypes. This is necessary for large-scale experiments with a large number of genotypes from different species that have widely different rates of growth.

The second objective was to determine the most informative traits indicating salinity tolerance in Galapagos tomato seedlings while highlighting the most tolerant accessions and their potential underlying mechanisms for salinity tolerance. It is widely known that salt stress affects many aspects of plant growth, such as biomass production, yield, photosynthesis and leaf metabolites (Chinnusamy *et al.*, 2006; Munns and Tester, 2008; Munns and Gilliham, 2015; Negrão *et al.*, 2017). Hence, there are many traits that could be recorded and analyzed to accurately assess the salinity tolerance of a plant. In this study, it was found that many of the traits were highly and significantly correlated, so the salinity tolerance could be assessed by focusing attention on only a few representative traits. From the highly-correlated leaf-related traits, leaf area was chosen as it was previously reported to be an important trait for salinity tolerance in tomato (Cuartero and Fernández-Muñoz, 1998; Dogan *et al.*, 2010). From the highly-correlated plant growth-related traits, the total fresh mass was chosen since it describes the increase in biomass and water retention at the whole plant level. Other traits, which did not correlate as strongly with the leaf- or growth-related traits, such as, root and stem length, leaf number, leaf elongation, leaf succulence, and Na and K concentrations in root and leaf, were included in further analyses. Leaf K was the only trait which negatively correlated with multiple traits. In general, the two Galapagos tomato species, *S. cheesmaniae* and *S. galapagense*, did not differentiate in PCA. This indicated high phenotypic variability within both species that was greater than differences between the two species, which are clear at the genetic level (Pailles *et al.*, 2017).

The PCA revealed that the selected traits contributing most to describe the variation across accessions were total fresh mass, leaf area, leaf succulence, and leaf K concentration. Plants with greater fresh mass and leaf area shared a tendency to have more leaves, longer stems and roots, and higher root K concentration, whereas leaf succulence, leaf elongation, K and Na concentrations in leaf, and root Na concentration tendencies differed greatly. The presence of similar tendencies in the two species suggests that salinity-tolerance related traits are conserved across these two species.

Total plant fresh mass, which is a measure of growth maintenance during salt stress, was the trait driving most of the variation across accessions. Growth maintenance has been widely acknowledged to be a good estimate of salinity tolerance (Genc *et al.*, 2007; Negrão *et al.*, 2017), especially at the seedling stage, since it is not possible in young plants to measure any commercially relevant traits such as yield. Thus, the genotypic variability for salinity tolerance was assessed in this study, based on the maintenance of growth under saline conditions relative to control conditions. This assumption was further confirmed by K-means clustering analysis, where the plant fresh mass of salt-treated plants relative to control plants was identified as the main trait driving the accessions clustering. The accessions were categorized into two groups with high or low salinity tolerance, based on the relative fresh mass. Within these groups, it was tested whether Galapagos tomato accessions showed co-varying groups of tolerance traits that are consistent across accessions, to categorize them as traits of influence in salinity tolerance. However, remarkable trait diversity was found within the highly-tolerant group, which suggests the presence of different mechanisms of salinity tolerance between accessions within the species, consistent with previous reports by Rush and Epstein (1981) and Cuartero et al. (1992).

Maintenance of growth, defined by an increase in mass, is one of the most important mechanisms contributing to salinity tolerance. According to Munns and Termaat (1986), leaf growth is more affected by salinity than root growth. In this study, leaf area was also observed to be an important trait contributing to plant mass. However, the physiological mechanisms underlying leaf growth inhibition under salt stress are not fully understood (Neves-Piestun and Bernstein, 2001). The current results showed a strong and significant correlation between leaf area and leaf elongation, followed by leaf area and leaf number. These found correlations could provide insights into the mechanisms by which salt stress affects leaf growth.

K concentration was found to be generally lower in all the high-tolerance accessions when compared to the low-tolerance accessions, especially in leaves. However, one group of *S. cheesmaniae* tolerant accessions showed higher levels of potassium in the root and leaf samples. Percey et al. (2016) reported that reduction in K^+^ efflux in halophytes is linked to reduced H^+^ efflux, which saves energy, allowing more resources to be redirected for plant growth. Therefore, the ability to maintain K^+^ uptake and a high K^+^:Na^+^ ratio under salt stress can be an important mechanism of salinity tolerance (Chen *et al.*, 2007; Shabala and Cuin, 2008). K^+^ deficiency in plants can impair photosynthesis (Cakmak, 2005), as well as many other aspects of cellular function, such as protein synthesis (Flowers and Dalmond, 1992). The four most salt tolerant accessions of *S. cheesmaniae* and the two most salt tolerant accessions of *S. galapagense* showed high K in their roots, which could indicate that they are good at maintaining K uptake under salt stress. In some accessions, higher K in the roots seems to go hand in hand with low K in the leaves, which could be explained by higher K^+^ re-translocation, useful to assist NO_3_^−^ uptake and distribution (Taleisnik and Grunherg 1994), or lower K^+^ translocation from roots to shoot. In addition, bigger leaves appeared to have a lower K concentration compared to smaller leaves, which could be due to a dilution effect, e.g. having a similar amount of K to that of the smaller leaves but more water content (Jarrell and Beverly, 1981).

Increase in leaf succulence (measured as water per unit leaf area), a strategy to reduce salt concentrations in photosynthetic tissues (Han *et al.*, 2013), is another known mechanism of salinity tolerance in some plants, including tomato (Cuartero and Fernández-Muñoz, 1998). The hierarchical clustering of accessions and traits showed that both *S. cheesmaniae* and *S. galapagense* accessions each formed a cluster of accessions with increased leaf succulence and low leaf Na concentrations. This might be caused by the succulence increasing cell size, thereby diluting the salt without increasing the leaf (Munns *et al.*, 2016).

The accumulation of Na^+^ relative to biomass can also be an indicator of salinity tolerance. However, Na^+^ is toxic when it accumulates in the cell cytosol, resulting in ionic disequilibrium (Hanin *et al.*, 2016). Additionally, Na^+^ reduces the availability of K^+^ binding sites for important metabolic processes in the cytoplasm (Wei *et al.*, 2017). For the plant to protect itself when exposed to salt stress, it has to either limit the entry of Na^+^ through the roots, or to control Na^+^ concentration and distribution once inside (Tester and Davenport, 2003; Hanin *et al.*, 2016). The Na^+^ that enters the root cells is extruded from the cytoplasm into the apoplastic space and compartmentalized into the vacuole (Maggio *et al.*, 2007). This process is called tissue tolerance (Munns *et al.*, 2016). One cluster from each species included tolerant accessions with high Na concentration in their leaves, which suggests significant levels of tissue tolerance (Munns *et al.* 2016). The S*. cheesmaniae* cluster appeared to respond to high Na accumulation in the leaves by growing more leaves, while the *S. galapagense* cluster appeared to increase leaf succulence.

In conclusion, the seedling screen study allowed characterization of the responses of 67 Galapagos tomato accessions to salt stress. Individual accessions were classified based on the phenotypic traits contributing to salinity tolerance. Interestingly, it was observed that individual salt tolerant accessions from the Galapagos Islands use different mechanisms to maintain their growth at the seedling stage under saline conditions. The different combinations of characteristics found across all the studied accessions, while maintaining a good relative fresh mass, indicate that the Galapagos tomatoes are naturally diverse and have different mechanisms to tolerate high salinity. In terms of growth maintenance under stress, the accessions LA0317, LA1449 and LA1403 displayed exceptional salinity tolerance at the seedling stage. However, to assess which mechanisms are the most effective for salinity tolerance, the tolerant accessions should be studied further at later growth stages, such as the reproductive stage, to evaluate the effect of salinity on yield. Additionally, trials to evaluate their performance under field conditions are recommended. Dissecting the genetic basis of salinity tolerance mechanisms through a genetic characterization and/or transcriptomic approach, together with our results, would facilitate the selection of useful accessions as genetic sources for breeding salinity tolerance traits into commercial tomatoes.

## Acknowledgements

We thank Igor Silva and Derek Burgess for assisting with logistics throughout the project, and Shireen Hammoud for assisting with sample collection and processing. We are also grateful to Muppala Reddy, Marina Khashat and Gomerito Sagun (KAUST greenhouse) for providing the experimental facilities and technical support. Significant text and content editing input from Neelima Sinha (UC Davis) is also gratefully acknowledged. The research reported in this publication was supported by funding from King Abdullah University of Science and Technology (KAUST). The authors have no conflicts of interest to declare.

